# Neural mechanisms of musical syntax and tonality, and the effect of musicianship

**DOI:** 10.1101/2022.10.14.512259

**Authors:** Lei Jiang, Ruiqing Zhang, Lily Tao, Yuxin Zhang, Yongdi Zhou, Qing Cai

## Abstract

The neural basis for the processing of musical syntax has previously been examined almost exclusively in classical tonal music, which is characterized by strictly organized hierarchical structure. The present study investigated the neural mechanisms for processing musical syntax across genres varying in tonality - classical, impressionist, and atonal music - and, in addition, examined how musicianship modulates such processing. Results showed that, first, the dorsal stream, including bilateral inferior frontal gyrus and superior temporal gyrus, plays a key role in the perception of tonality. Second, right fronto-temporal regions were crucial in allowing musicians to outperform non-musicians in musical syntactic processing; musicians also benefit from a cortical-subcortical network including pallidum and cerebellum, suggesting more auditory–motor interaction in musicians than in non-musicians. Third, left pars triangularis carries out on-line computations independently of tonality and musicianship, whereas right pars triangularis is sensitive to tonality and partly dependent on musicianship. Finally, unlike tonal music, processing of atonal music could not be differentiated from that of scrambled notes, both behaviorally and neurally, even among musicians. The present study highlights the importance of studying varying music genres and experience levels, and provides a better understanding of musical syntax and tonality processing and how such processing is modulated by music experience.

## 1 Introduction

Throughout the history of humanity, music has been a key component in social and cultural interactions. How people communicate with music, namely how listeners perceive music syntax has been the subject of investigation in neuroscience. Some have suggested parallels between music processing and language processing. Currently, however, the neural mechanisms of tonal music perception are still uncertain. Some evidence has been provided by studies on Western classical music. The organization of pitches or chords in classical harmonic musical sequence tends to begin with the main tone or chord, and usually returns to the main tone or chord at the end. Other genres of music involve different structures, and may, thus, entail different processing mechanisms to classical music.

Animal studies have shown that, in marmosets, harmonic template neurons sensitive to spectral regularity of harmonic complex sounds are distributed across the primary auditory cortex and the neighboring primary-like rostral area (Feng & Wang, 2017). In humans, widely distributed frontal and temporal regions have been involved in the precessing of classical music. Among these regions, the left inferior frontal gyrus (IFG) has been suggested to be the most important site offering computational resources for both linguistic and musical syntax (Patel, 2003, Patel et al., 2008; Kunert et al., 2015). Electrophysiological studies have suggested that patients with lesions in left IFG show abnormal musical syntax processing and impaired behavioral performance in the processing of irregular chord sequences, and that left IFG is the key region for the processing of syntax in a domain-general way (Sammler, Koelsch, &Friederici, 2011; Patel et al., 2008). Furthermore, music processing, like language processing, may also involve shared dorsal and ventral neural networks, underlying structure and meaning processing respectively (Koelsch & Siebel, 2005; Musso et al., 2015). The dorsal stream – including IFG, anterior superior temporal gyrus (STG) and ventrolateral premotor cortex (PMC) – processes harmonic relations and structural irregularities, predicts short-term upcoming harmonic sequences (Koelsch & Siebel, 2005), and is involved independently of the type of musical stimuli (Tillmann et al., 2006). The left IFG further connects to inferior parietal cortex and middle temporal lobe through dorsal and ventral long association tracts (Musso et al., 2015).

Although previous studies have provided a good basis for the understanding of music processing, so far almost all neuroscientific studies on music exclusively used Western classical music. Classical Western music is characterized by strictly organized hierarchical structure, which may not be the case across other music genres. It is important, therefore, to examine a variety of music genres to provide a complete and unbiased picture (see also Brattico et al., 2013). Let us take a closer look at two other music genres: impressionist music and atonal music. Representative compositions of impressionism are partial to the diatonic scale. Impressionist musicians such as Debussy divides an octave into six major second intervals of three kinds– major second, major third, and tritone (Day-O’Connell, 2009). Atonal music exploits a composition technique without the tonic center and functional relationship among notes or chords. For example, in “A Survivor of Warsaw”, a representative atonal piece written by Schoenberg, the twelve semitones are functionally equal, making it distinct from the major-minor system. Moreover, the size distribution of intervals in the scale of tonal music is generally between one and three half-tones, and the grading progress is the main composition of the melody lines.

In short, the diatonic scale in impressionist music and the combination of 12 equal half-tones in atonal music both break the structural rules of classical music, either partially or completely. The asymmetry of the scale, the limitation of sound levels, and the size distribution of intervals within the scale are some important factors that differentiate tonal, impressionist, and atonal music in music theory. According to the literature on music processing, if the interval relationship to the tonal center (i.e. pitch-center relationship) disappears, the musical grammar would be disrupted and listeners could feel weary (Lerdahl & Jackendoff, 1983). If this is the case for atonal music, we should expect that the neural networks underlying the processing of the regularities of pitch relationships and structure-based prediction to also work differently.

A further question is whether such neural activation is exclusively decided by the physical features of musical stimuli, which is identical for all listeners; or if it rather reflects how the music is perceived by individuals and, therefore, interacts with listeners’ music experience and preference. For example, for a non-trained listener, music may simply be a series of notes and beats, sometimes even a nuisance to the ear. For the romantic musician, in contrast, music can communicate just as well, or even better than language. In other words, training and experience matters. Previous findings have shown that the early right anterior negativity (ERAN) ERP component is sensitive to music training (Koelsch et al., 2002b). A recent study further showed that, in musicians, right IFG, as well as right posterior STG, superior temporal sulcus (STS), and cerebellum are involved in the processing of musical structures, with resting state activity in right IFG positively correlated with that in posterior STG and left Heschl’s gyrus (Bianco et al., 2016). However, only musicians were tested in that study, so it remains unclear how music experience modulates music processing and whether this process interacts with tonality.

The present study aimed to investigate the neural mechanisms underlying the processing of musical syntax, as well as the impact of tonality and expertise on such processing. To achieve this purpose, we included music genres that varied in tonality. Specifically, extending from previous studies on classical tonal music, we also examined impressionist music (relatively decreased tonality) and atonal music (no tonality). A second aim of the present study was to investigate how musicianship modulates musical structure processing, and how it interacts with different music genres – that is, whether music experience affects brain networks underlying music tonal syntactic processing.

## 2 Materials and Methods

### 2.1 Participants

Thirty-six healthy native Chinese speakers with normal hearing, recruited from East China Normal University or Shanghai Conservatory of Music, took part in this study. All participants were right-handed, confirmed using Edinburgh Handedness Inventory (Oldfield, 1971). Written informed consent was obtained from each participant, and the protocol of the present study was approved by the Committee on Human Research Protection at East China Normal University. All participants were paid for their participation.

Musicianship was determined using Music Experience Questionnaire. Half of the participants (n=18) were musicians (22.4 ±2.1 years, 16 females) who majored in instrumental (17) or vocal (1) performance, and were immersed in a classical music environment for on average 3.3 (±2.4) hours per day; five of them reported having absolute pitch. They had on average 13.0 years of formal music training (± 3.2, range 8 to 17 years), with an average age of onset of 5.5 years (± 1.3, range 3 to 8 years).

The other half of the participants (n=18) were non-musicians (21.3 ± 3.3years, 13 females), who reported no prior experience in music training except one with a one-year experience in learning accordion and two with limited experience of playing piano or keyboard at young ages (these three participants took part in the study given their limited music experience and no music training in the last ten years, but their data were excluded in further analysis).

### 2.2 Materials

There were three experimental conditions, that is, three genres of music – classical/tonal, impressionist/pantonal, atonal – and three control conditions – their respective scrambled versions. In order to inspect more global and salient violations of tonal syntax, we adopted a method used in Levitin and Menon (2003), in which scrambled versions of musical pieces were included as baseline conditions to disrupt the musical structure, in other words the overall relationship between adjacent notes.

Each of the three experimental conditions contained 40 phrases, selected from representative Western composers’ masterpieces, as listed in Table 1. The phrases were reconstructed using Sibelius software to be synchronous, to have a similar number of notes (32±2 notes), and similar intensity. Only the relative positions of the notes, or the internal organizational structure of the phrase, was preserved. By doing so, the low-level acoustic features such as tempo, loudness, and timbre were balanced across music genres and leave the music syntax intact. The mean duration of the phrases was 6.2 (±0.4) s. Scrambled versions were made by shuffling all of the notes within each of the original phrases, so that the relative pitch of adjacent notes was disrupted. The scrambled phrases were then rated by three professional musicians independently to ensure that the inner original organizational structures had been destroyed while the same notes were kept. To increase relative loudness of pitch in the noisy scanner environment, dynamic range compression was applied on all the pieces using the compressor effect of Audacity (Farbood, 2015).

**Table 1.**
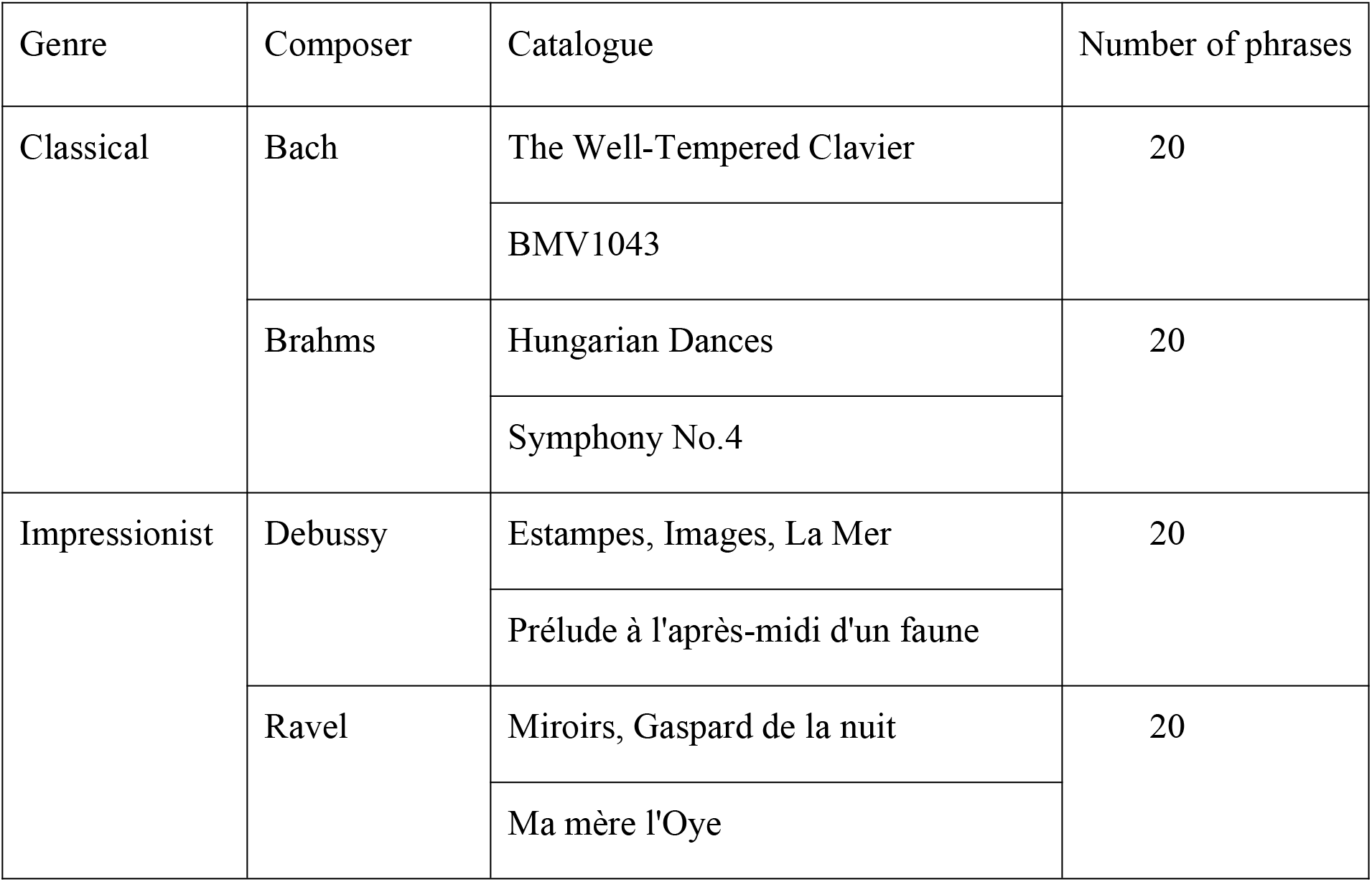
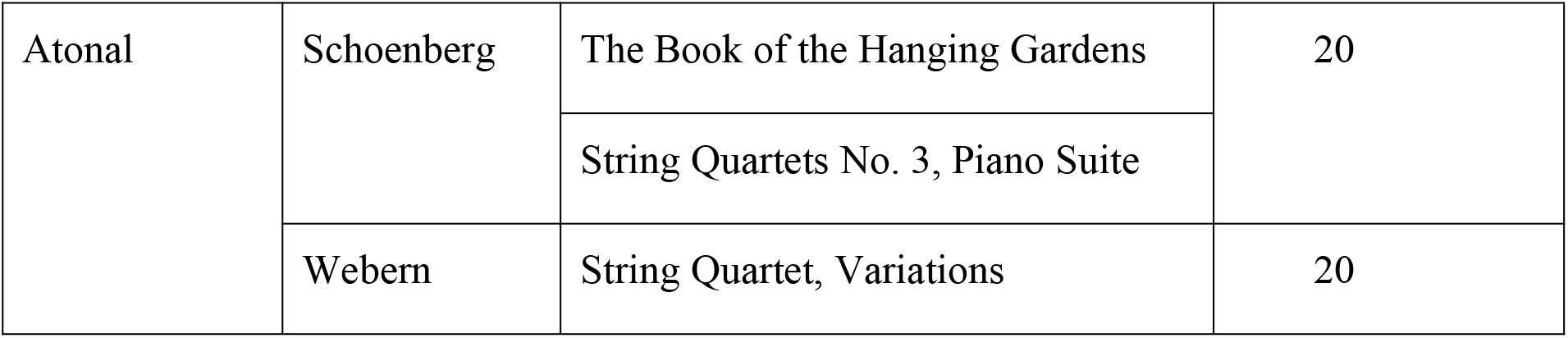
List of sources for the three genres of musical materials.

| Genre | Composer | Catalogue | Number of phrases |
| --- | --- | --- | --- |
| Classical | Bach | The Well-Tempered Clavier | 20 |
|  |  | BMV1043 |  |
|  | Brahms | Hungarian Dances | 20 |
|  |  | Symphony No.4 |  |
| Impressionist | Debussy | Estampes, Images, La Mer | 20 |
|  |  | Prélude à l'après-midi d'un faune |  |
|  | Ravel | Miroirs, Gaspard de la nuit | 20 |
|  |  | Ma mère l'Oye |  |
| Tonality decides how much we can appreciate the music. |  |  |  |
| Atonal | Schoenberg | The Book of the Hanging Gardens | 20 |
|  |  | String Quartets No. 3, Piano Suite |  |
|  | Webern | String Quartet, Variations | 20 |

In addition to these stimuli, a 250-Hz pure tone (660 ms duration) was used as probe stimulus. Five such trials were included, inserted evenly between other trials, within each scanning session/run, to ensure that participants were attending to the task.

### 2.3 Procedure

During fMRI scanning, participants were required to listen carefully to each phrase presented (they were not informed that some are original and others are scrambled), and to press a button with their right index finger when they hear the pure tone (which had been presented to them outside before scanning). The same task was performed twice in the scanner. Each session/run contained 125 trials: 20 trials for each of the six conditions plus five pure-tone probe-detection trials. Each session/run started with a fixation of 10 s, and then all trials were presented in a random order. Between each phrase, a 2-4-6 s blank interval was presented (see Figure 1A). Stimuli were presented using E-Prime 2.0 software.

**Figure 1.**
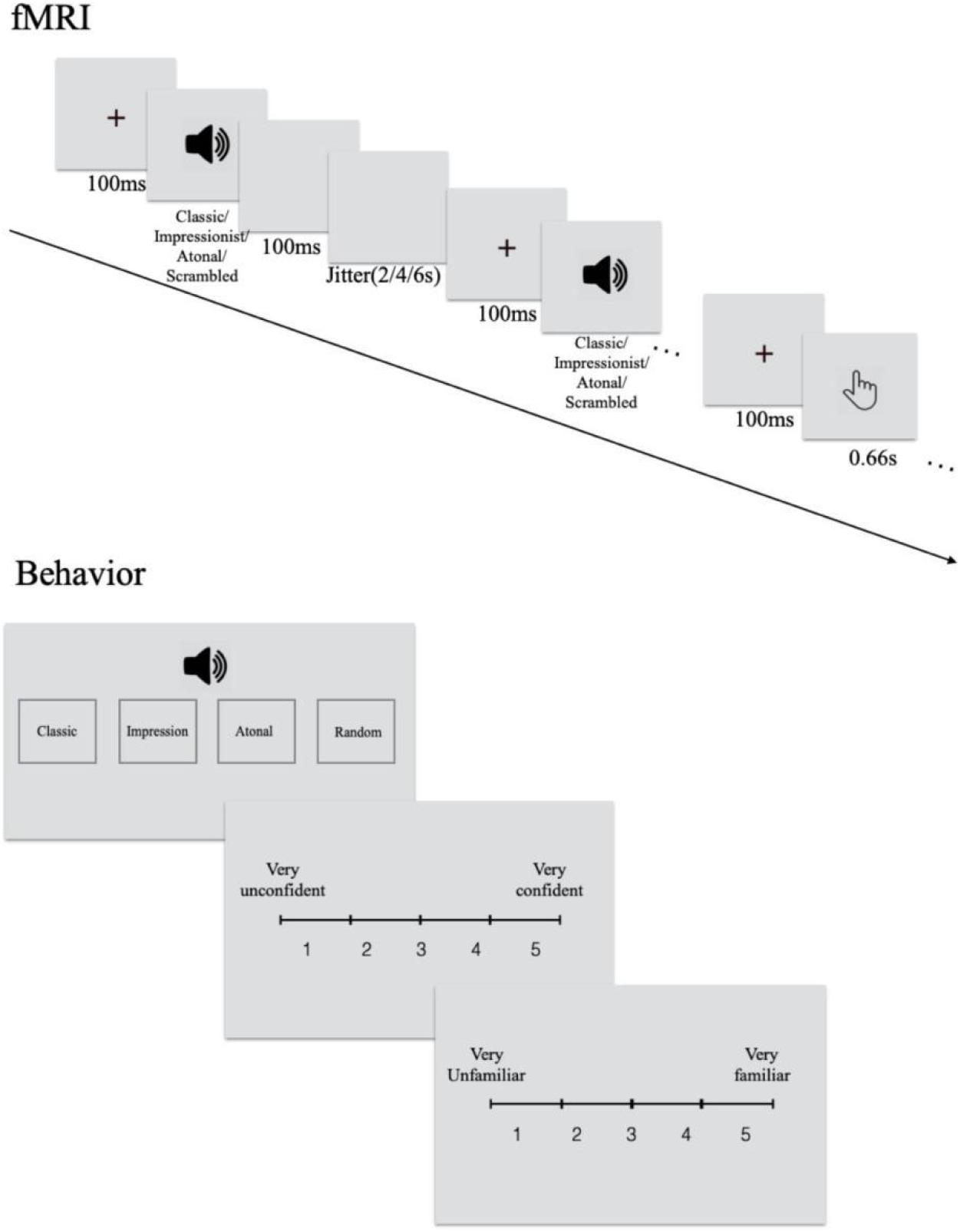
Experimental procedure for (A) fMRI and (B) behavioral tasks.

After scanning, participants listened to all phrases again, classified each piece into four categories (classical/tonal, impressionist, atonal music, and random notes), rated the level of confidence in his/their decision (from 1 = least confident to 5 = most confident), and familiarity with the phrase (from 1 = least familiar to 5 = most familiar; see Figure 1B).

### 2.4 Data Acquisition

Whole-brain images were collected on a 3T Siemens Trio MR scanner, with a 32-channel head coil. First, an anatomical image was obtained using a T1-weighted MPRAGE sequence (TR = 2530 ms, TE = 2.34 ms, image matrix = 256 * 256, FoV = 256 mm, flip angle = 7°, voxel size = 1*1*1mm, 192 slices). Functional MRI images were acquired using a T2*-weighted gradient-echo EPI sequence covering the whole brain (TR= 2400 ms, TE = 30 ms, image matrix = 64*64, FoV = 192 mm, flip angle = 81°, voxel size = 3*3*3mm, slice thickness = 3mm, 40 slices, interleaved acquisition). Stabilization cushions were used to minimize head motion, and ear plugs were worn by participants to reduce noise from the scanner during operation. Auditory stimuli were presented using RT-300 (Resonance Technology, Canada). Behavioral data were collected outside the MRI environment after scanning.

### 2.5 Behavioral Data Analysis

Two-way mixed design ANOVA with Tukey’s HSD comparison tests were performed separately for the genre classification, confidence rating, and familiarity rating, with group (musician, non-musician) and musical syntax (classical, impressionist, atonal, random notes) as independent factors. Data from two musicians were excluded because their accuracy for “random notes” were outliers (0% and 2.5%). Note that for each participant and each genre, familiarity score was calculated based on ratings for all phrases, and confidence score only took into account the correctly classified trials.

### 2.6 Functional Imaging Data Analysis

Functional MRI data preprocessing and statistical analysis was carried out using SPM8 (www.fil.ion.ucl.ac.uk/spm). After slice-timing correction, the functional images were realigned for headmotion correction. The functional and co-registered anatomical images were spatially normalized to MNI space, and then smoothed using a Gaussian kernel with full width at half maximum (FWHM) of 5 mm. Head movements were checked for each subject using Artifact Detection Tools (ART; www.nitrc.org/projects/artifact_detect) package. Time points (scans/volumes) with motion outliers (≥2 mm) or outliers in global signal intensity (≥5 SD) were recorded for nine participants.

Data from each participant were then analyzed using a general linear model (GLM), with three music genre conditions (classical, impressionist, and atonal), three scrambled conditions, and the probe condition. Head movement parameters were included for each participant as regressors, and the above mentioned time points with motion or intensity outliers were omitted by including a single regressor for each in GLM. Familiarity scores from participants’ behavioral ratings were included as parametric modulators for each condition to dissociate familiarity effects from main effects.

We first examined whether there were significant differences between any two scrambled conditions (out of the three scrambled conditions) using a 2 (group) * 3 (condition) flexible factorial model at the group level. Given no significant main effect or interaction was found for the scrambled conditions, the three scrambled conditions were combined into one, referred to as the random notes condition (matching the music genre classification in behavioral analysis). A 2 (group) * 4 (musical syntax: classical, impressionist, atonal, random notes) flexible factorial model was used in further analysis.

Given previous discoveries on the functional role of bilateral IFG in music processing, bilateral IFG (pars triangularis and pars opercularis) anatomical ROIs were selected from MarsBaR AAL ROIs. Percent signal change relative to global brain signal was computed using MarsBar, to further investigate how the brain reacted to different music genres in musicians and non-musicians.

To further derive the synchronous function of cortical regions that processed different music genres in musicians and non-musicians separately, informational connectivity analysis (Coutanche & Thompson-Schill, 2013) was conducted. The whole brain was segmented into 116 regions of interest (ROIs) based on Automated Anatomical Labeling 116 (AAL116) template (Tzourio-Mazoyer et al., 2002; Schmahmann et al., 1999). Four ROIs were excluded in further analysis because they have not been fully covered in certain participants while scanning. For each ROI, a representational dissimilarity matrix (RDM) of all 240 musical trials was computed based on ß values extracted from all voxels for each participant. Then, for each ROI pair, the correlation coefficient was calculated between the two RDMs of the ROI pair and then transformed to fisher’s z values indicating representational similarity of general musical sentences processing between brain regions. After that, the correlation analysis was then performed separately for musicians and non-musicians to investigate the relationship between the z values of each region pair and the behavioral overall genre classification accuracy (representing each participant’s general musical genre sensitivity). Informational connectivity analysis allows us to inspect the highly stimuli-dependence neural processing between brain regions, which offers a higher-order explanation than univariate analysis.

## 3 Results

### 3.1 Behavioral Results

ANOVA on classification accuracy showed a main effect of group, F(1,136)=25.058, p<0.001, a main effect of musical syntax F(3,136)=19.135, p<0.001, and an interaction between group and musical syntax (F(3,136)=4.837, p<0.01). Tukey’s HSD post-hoc test indicated that classical music and impressionist music were easier to identify than atonal music (HSD =24.118, p<0.001, HSD =20.881, p<0.001, respectively classical and impressionist) and random notes (HSD =15.662, p<0.001, HSD =12.425, p<0.01, respectively for classical and impressionist). The musician group classified classical and impressionist music better than atonal music (HSD =23.594, p<0.001, HSD=28.292, p<0.001, respectively for classical and impressionist) and random notes (HSD=22.552, p<0.001, HSD=27.25 p<0.001, respectively for classical and impressionist). The non-musician group was found to have better knowledge only of classical compared to atonal genre (HSD=24.583, p<0.001). Within music genres, a significant group difference was only found for impressionist music classification, with musicians outperforming non-musicians (HSD=28.083, p <0.001; see Figure 2A).

**Figure 2.**
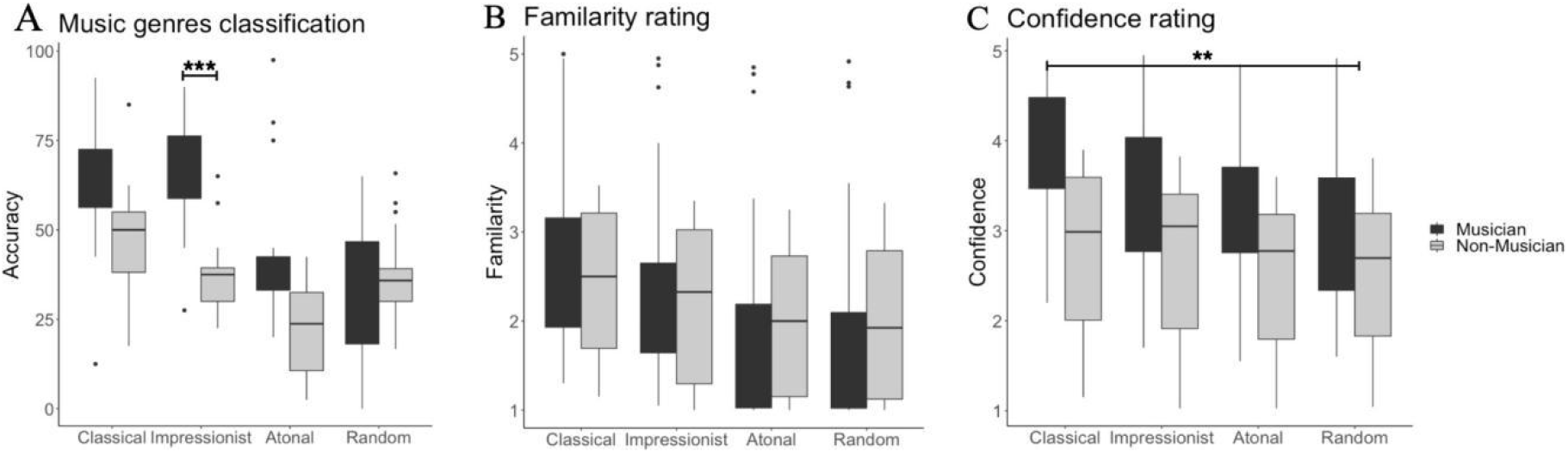
Behavioral results for musicians and non-musicians for (A) percentage correct genre classification, (B) familiarity ratings (1, least familiar∼5, most familiar) in musicians and non-musicians, and (C) confidence ratings (1, least confident∼5, most confident).

For familiarity ratings, ANOVA showed only a significant main effect of musical syntax, F(3,140)=21.91, p <0.001. Post-hoc tests showed that classical musical phrases were rated as significantly more familiar than atonal musical phrases (HSD=0.56, p<0.05), and significantly more familiar than random notes (HSD=0.638, p <0.01; see Figure 2B).

For confidence ratings, ANOVA showed significant main effects of groups, F(3,142)=9.079, p<0.01, and of musical syntax, F(3,140)=23.657, p<0.001. Musicians were overall more confident than non-musicians in their genre classifications (HSD=0.896, p<0.001). Confidence was significantly higher when classifying classical music comparing to atonal music (HSD=0.727, p<0.01) and random notes (HSD=0.676, p<0.01; see Figure 2C).

### 3.2 Functional Imaging Results

The group-level factorial analysis showed a significant interaction between group and musical syntax, which involved activation in right postcentral areas, left supplementary motor area (SMA), left middle temporal gyrus (MTG), left hippocampus, and bilateral superior frontal gyrus (SFG). The main effect of musical syntax was observed in bilateral superior temporal regions, bilateral IFG pars triangularis extending to left insula, bilateral superior medial frontal areas, bilateral precentral gyrus, right SFG, right middle frontal gyrus (MFG), left angular gyrus, right supramarginal gyrus, left SMA, and bilateral cerebellum. The main effect of group was observed in bilateral cerebellum, bilateral precentral gyrus, right SFG, right superior temporal pole, bilateral inferior temporal gyrus, left amygdala, right STG, and bilateral IFG pars opercularis (all p’s<0.001, alphasim corrected; see Table 2).

**Table 2.**
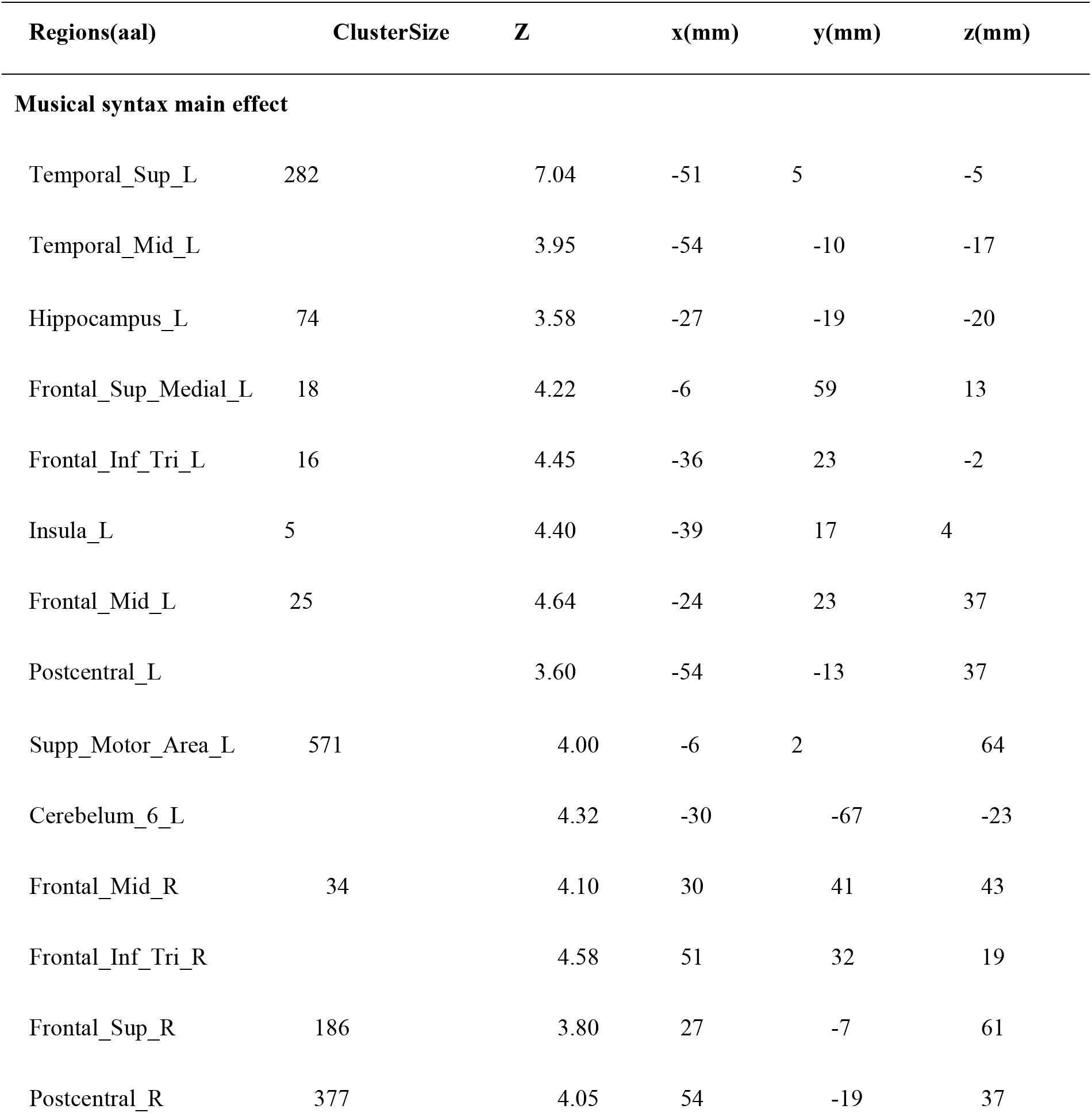
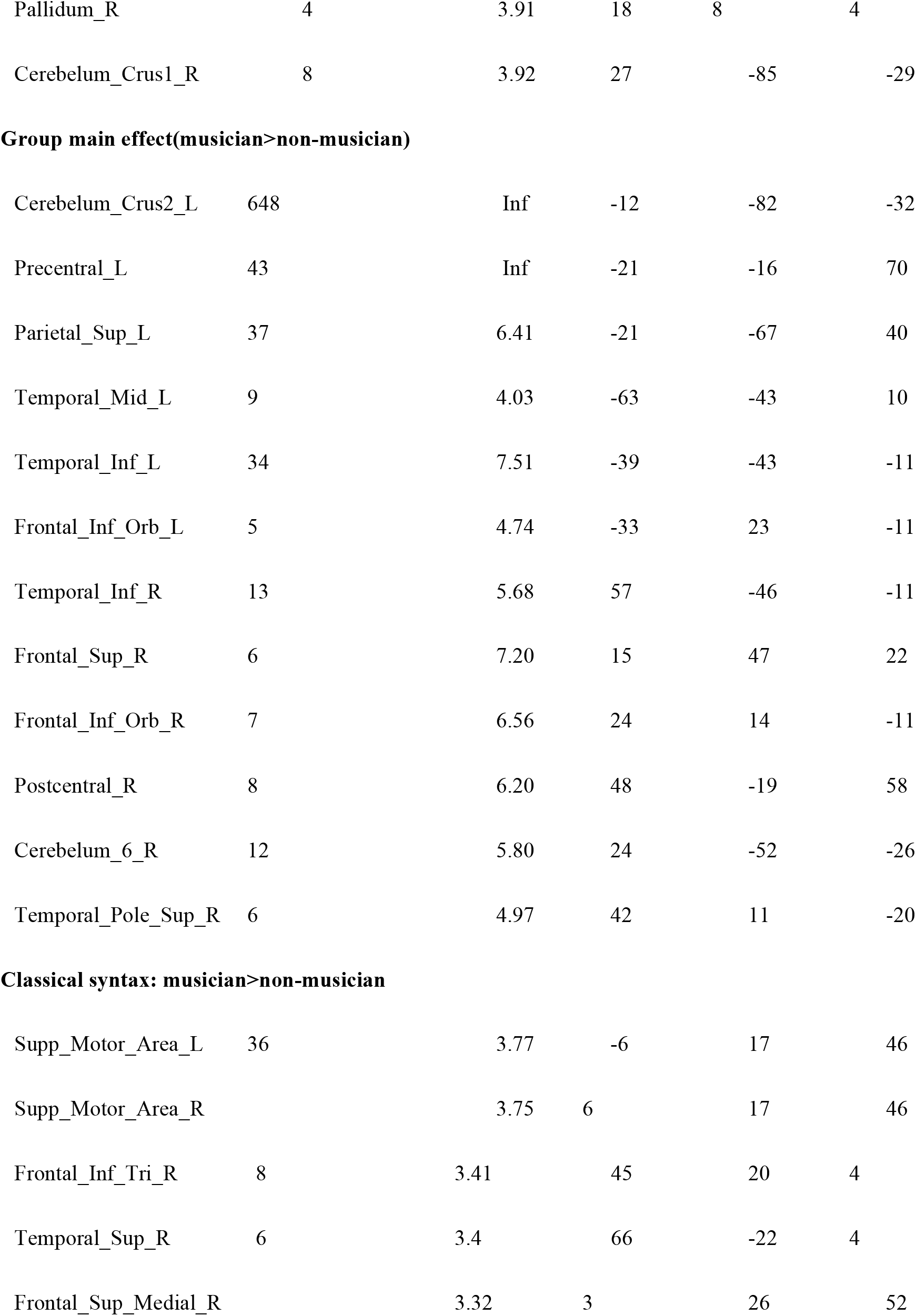
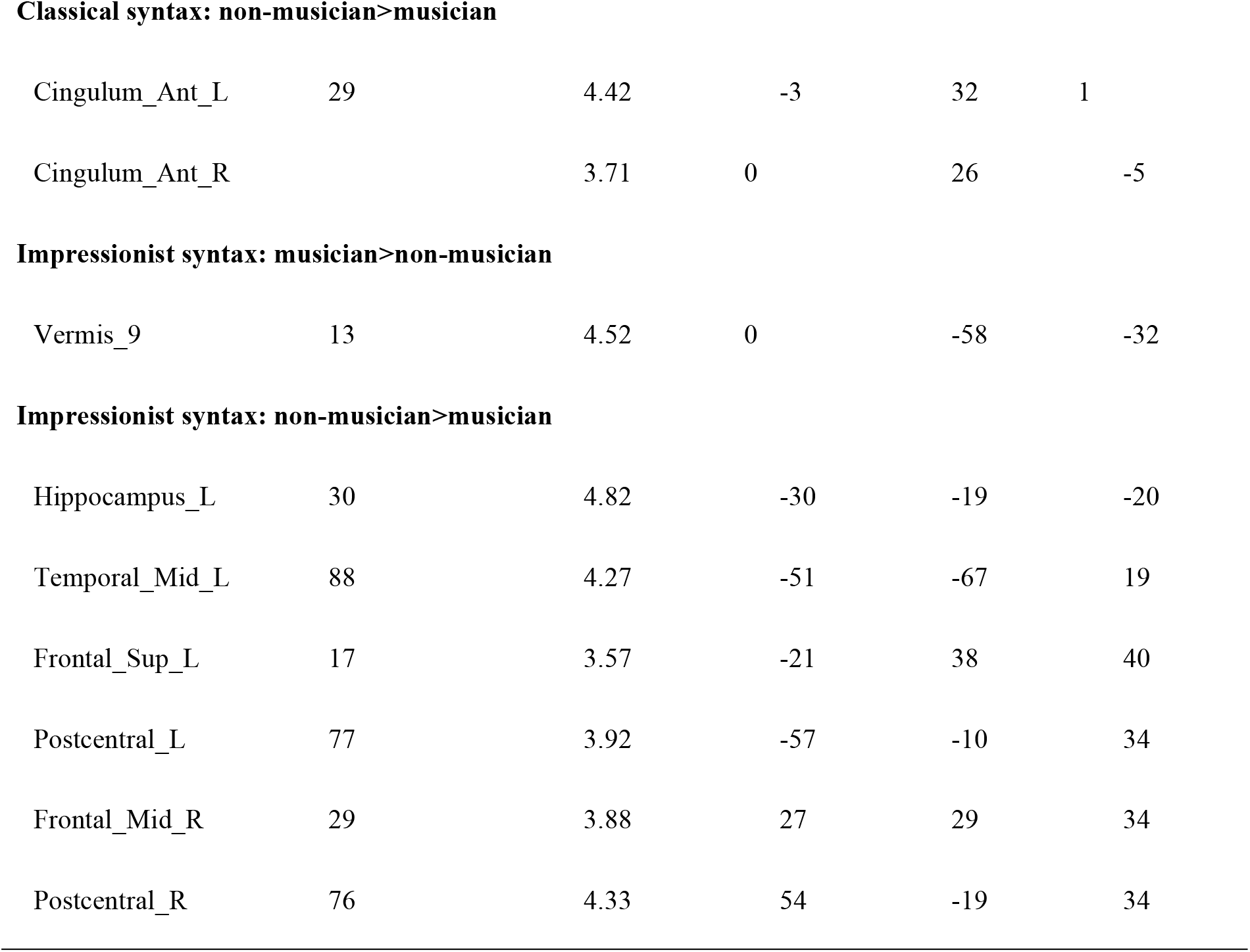
Activation results of main effects of musical syntax and group, and simple effects of musicianship on classical and impressionist music processing (all alphasim corrected at p<0.001).

| Regions(aal) | ClusterSize | Z | x(mm) | y(mm) | z(mm) |
| --- | --- | --- | --- | --- | --- |
| **Musical syntax main effect** |  |  |  |  |  |
| Temporal_Sup_L | 282 | 7.04 | -51 | 5 | -5 |
| Temporal_Mid_L |  | 3.95 | -54 | -10 | -17 |
| Hippocampus_L | 74 | 3.58 | -27 | -19 | -20 |
| Frontal_Sup_Medial_L | 18 | 4.22 | -6 | 59 | 13 |
| Frontal_Inf_Tri_L | 16 | 4.45 | -36 | 23 | -2 |
| Insula_L | 5 | 4.40 | -39 | 17 | 4 |
| Frontal_Mid_L | 25 | 4.64 | -24 | 23 | 37 |
| Postcentral_L |  | 3.60 | -54 | -13 | 37 |
| Supp_Motor_Area_L | 571 | 4.00 | -6 | 2 | 64 |
| Cerebelum_6_L |  | 4.32 | -30 | -67 | -23 |
| Frontal_Mid_R | 34 | 4.10 | 30 | 41 | 43 |
| Frontal_Inf_Tri_R |  | 4.58 | 51 | 32 | 19 |
| Frontal_Sup_R | 186 | 3.80 | 27 | -7 | 61 |
| Postcentral_R | 377 | 4.05 | 54 | -19 | 37 |
| Pallidum_R | 4 | 3.91 | 18 | 8 | 4 |
| Cerebelum_Crus1_R | 8 | 3.92 | 27 | -85 | -29 |
| Group main effect(musician>non-musician) |  |  |  |  |  |
| Cerebelum_Crus2_L | 648 | Inf | -12 | -82 | -32 |
| Precentral_L | 43 | Inf | -21 | -16 | 70 |
| Parietal_Sup_L | 37 | 6.41 | -21 | -67 | 40 |
| Temporal_Mid_L | 9 | 4.03 | -63 | -43 | 10 |
| Temporal_Inf_L | 34 | 7.51 | -39 | -43 | -11 |
| Frontal_Inf_Orb_L | 5 | 4.74 | -33 | 23 | -11 |
| Temporal_Inf_R | 13 | 5.68 | 57 | -46 | -11 |
| Frontal_Sup_R | 6 | 7.20 | 15 | 47 | 22 |
| Frontal_Inf_Orb_R | 7 | 6.56 | 24 | 14 | -11 |
| Postcentral_R | 8 | 6.20 | 48 | -19 | 58 |
| Cerebelum_6_R | 12 | 5.80 | 24 | -52 | -26 |
| Temporal_Pole_Sup_R | 6 | 4.97 | 42 | 11 | -20 |
| Classical syntax: musician>non-musician |  |  |  |  |  |
| Supp_Motor_Area_L | 36 | 3.77 | -6 | 17 | 46 |
| Supp_Motor_Area_R |  | 3.75 | 6 | 17 | 46 |
| Frontal_Inf_Tri_R | 8 | 3.41 | 45 | 20 | 4 |
| Temporal_Sup_R | 6 | 3.4 | 66 | -22 | 4 |
| Frontal_Sup_Medial_R |  | 3.32 | 3 | 26 | 52 |
| Classical syntax: non-musician>musician |  |  |  |  |  |
| Cingulum_Ant_L | 29 | 4.42 | -3 | 32 | 1 |
| Cingulum_Ant_R |  | 3.71 | 0 | 26 | -5 |
| Impressionist syntax: musician>non-musician |  |  |  |  |  |
| Vermis_9 | 13 | 4.52 | 0 | -58 | -32 |
| Impressionist syntax: non-musician>musician |  |  |  |  |  |
| Hippocampus_L | 30 | 4.82 | -30 | -19 | -20 |
| Temporal_Mid_L | 88 | 4.27 | -51 | -67 | 19 |
| Frontal_Sup_L | 17 | 3.57 | -21 | 38 | 40 |
| Postcentral_L | 77 | 3.92 | -57 | -10 | 34 |
| Frontal_Mid_R | 29 | 3.88 | 27 | 29 | 34 |
| Postcentral_R | 76 | 4.33 | 54 | -19 | 34 |

Overall, classical/tonal music (compared to random notes) involved significant activation in bilateral STG, left inferior frontal regions (including pars triangularis, pars opercularis, and pars orbitalis), right inferior frontal regions (including pars opercularis and insula), bilateral precentral gyrus, bilateral SMA, and bilateral cerebellum. Impressionist music (compared to random notes) involved significant activation in bilateral STG, right superior temporal pole, right MTG, left IFG pars opercularis and pars triangularis, right IFG pars triangularis, left supramarginal gyrus, right hippocampus, right precentral gyrus, right SMA, and left cerebellum. When contrasting classical over impressionist music processing, classical condition involved greater activation in right IFG pars opercularis and left insula compared to impressionist; the reverse contrast involved more right IFG pars triangularis, right precentral gyrus, bilateral STG, bilateral superior temporal pole, and left SMA. When comparing to atonal music, classical music showed more activation in bilateral STG and MTG, right IFG pars triangularis and pars opercularis, left IFG pars opercularis and insula, bilateral precentral gyrus, bilateral SMA, and bilateral cerebellum; impressionist music showed more activation in bilateral STG and MTG, left IFG pars triangularis and pars orbitalis, right IFG pars triangularis, bilateral SMA, bilateral putamen, and bilateral cerebellum. Atonal music involved more activation in bilateral MTG than classical music, with no areas showing greater activation compared to impressionist music (all *p’s* < 0.001, alphasim corrected; see Figure 3A-C).

**Figure 3.**
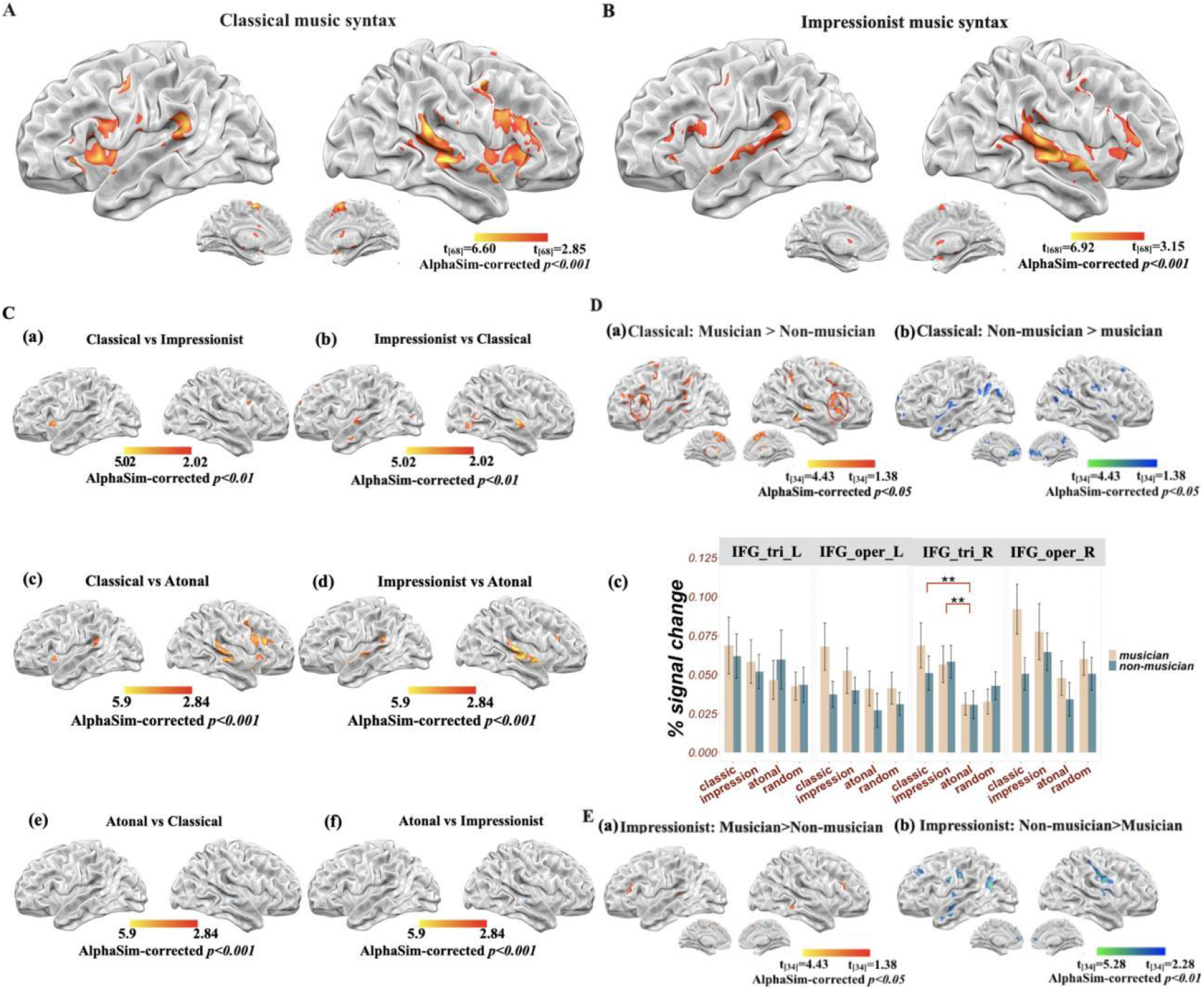
Brain activation for musical syntax processing (all results alphasim corrected at p<0.001, unless otherwise stated). (A) Classical music compared to random notes; (B) impressionist music compared to random notes; (C) Comparisons among musical genres: a) classical compared to impressionist music (alphasim corrected at p<0.01 for illustration); b) impressionist compared to classical music (alphasim corrected at p<0.01 for illustration); c) classical compared to atonal music; d) impressionist compared to atonal music; e) atonal compared to classical music; f) atonal compared to impressionist music; (D) difference between groups for classical music: a) musicians compared to non-musicians; b) non-musicians compared to musicians; c) percent signal change in left and right IFG; (E) differences between groups for impressionist music: a) musicians compared to non-musicians; b) non-musicians compared to musicians.

Simple effects were further analyzed using t-tests to investigate how the processing of musical structure was modulated by musicianship. For classical music processing, musicians showed greater activation in right STG, right IFG pars triangularis, right superior medial frontal gyrus, right inferior parietal gyrus, and bilateral SMA, whereas bilateral anterior cingulate cortex (ACC) were more activated in non-musicians (all *p’s* < 0.001, alphasim corrected; see Table 2).

When processing impressionist music, musicians showed more activation in left cerebellum (Vermis 9) compared to non-musicians; non-musicians showed more activation in bilateral hippocampal gyrus, bilateral postcentral gyrus, MTG, SFG, insula, precuneus, and middle occipital lobe in the left hemisphere (all *p’s* < 0.001, alphasim corrected; see Table 2).

Lastly, for atonal music, no significant differences were found between musicians and non-musicians (*p’s* < 0.001, alphasim corrected).

For the ROI analysis on bilateral IFG (see Figure 3D(c)), left pars triangularis showed no sigificant effects of music genre (*F*_(3)_ = 0.842, *p* = 0.472) or group (F_(1)_ = 0.000, *p* = 0.984), or their interaction (*F*_(1,3)_ = 0.218, *p* = 0.884). A significant musical syntax main effect (*F*_(1,3)_ = 0.048, *p* < 0.01) was found for right pars triangularis, specifically, both classical (t = 2.76, *p* < 0.01, 95% CI = [0.0112,0.0222]) and impressionist music (t = 3.28, *p* < 0.01, 95% CI = [0.0683,0.0745]) have greater signal change than atonal music. For both left and right pars opercularis, there were significant group differences (left: *F*_(1,3)_ = 4.61, *p* < 0.05; right: *F*_(1,3)_ = 4.65, *p* < 0.05), with percent signal change in musicians greater than in non-musicians (left: t = 2.15, *p* < 0.05, 95% CI = [0.0014,0.0322]; right: t = 2.18, *p* < 0.05, 95% CI = [0.0553,0.0459]).

Informational connectivity between right Heschl’s gyrus and right superior temporal pole was positively correlated with behavioral classification accuracy in musicians (*r* = 0.69, FDR corrected at *q* = 0.005); informational connectivity between right IFG pars orbitalis and left pallidum was also positively correlated with behavioral classification accuracy in musicians (*r* = 0.79, FDR corrected at *q* = 0.005). Informational connectivity between cerebellum (cerebellar vermis 7, VER7) and both left and right STG was negatively correlated with behavioral accuracy in non-musicians (left STG: *r* = −0.89, *q* = 0.0001; right STG: *r* = −0.74, *q* = 0.0001) (see Figure 4).

**Figure 4.**
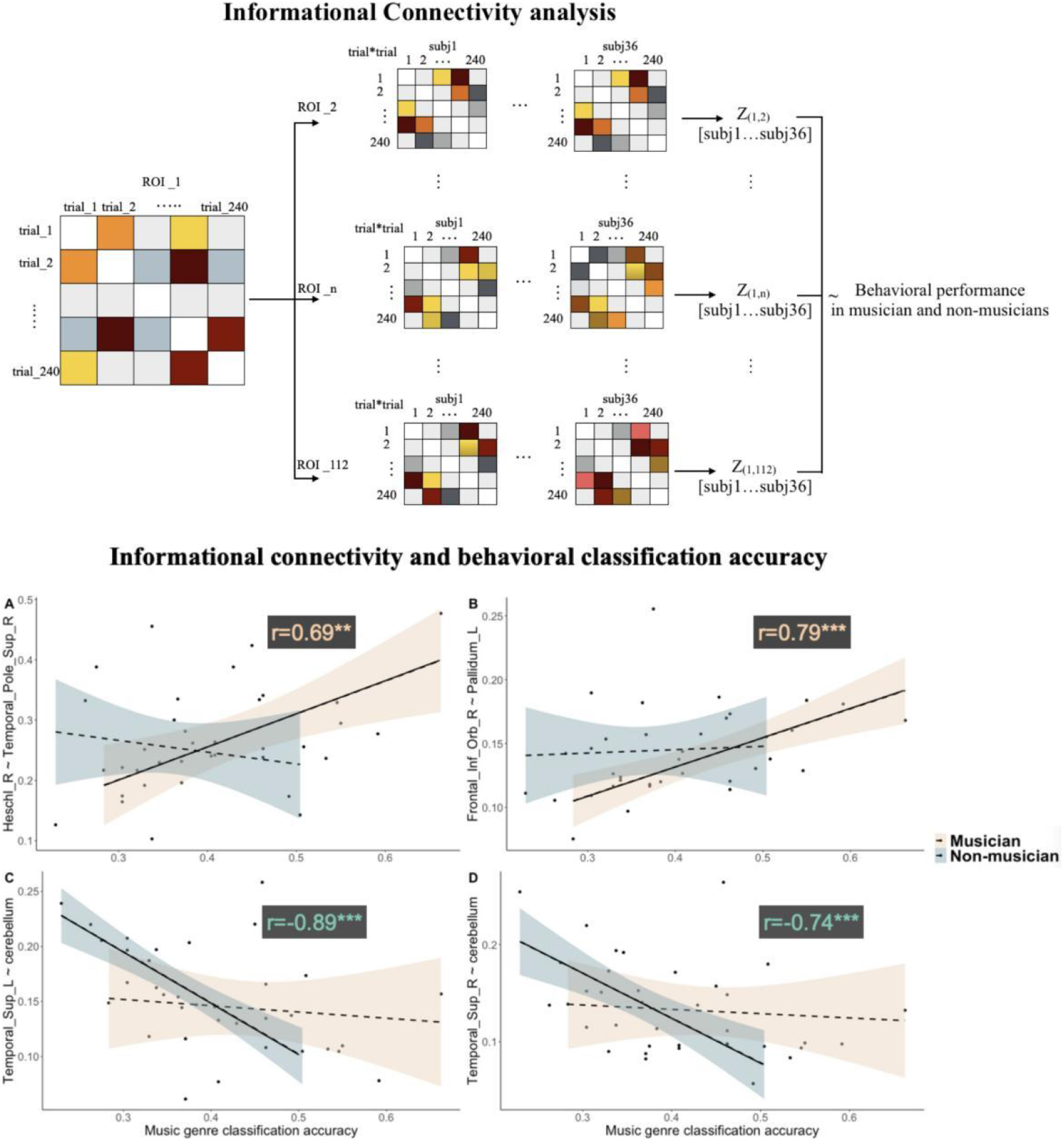
Upper panel: illustration for computing informational connectivities between ROIs for all the participants. Lower panel: correlations between informational connectivities and behavioral classification accuracy in musicians and non-musicians, for (A) connectivity between right Heschl’s gyrus and right superior temporal pole, (B) connectivity between right IFG pars orbitalis and left pallidum, (C) connectivity between the left superior temporal gyrus and cerebellum, and (D) connectivity between right superior temporal gyrus and cerebellum.

## 4 Discussion

The present study investigated the neural mechanisms underlying tonality and musical syntax processing, as well as the role of music training on such processing. Musicians and non-musicians listened to phrases from classical, impressionist, and atonal music genres inside an MRI scanner, and performed a classification task outside the scanner. The results elucidated the on-line processing mechanisms of musical syntax across different genres, and showed how musicianship impacted the neural response to different musical syntax.

### 4.1 Musical syntax, tonality, and musicianship

For overall processing of hierarchical structure in music, neural response was observed in bilateral temporal lobes, IFG, postcentral gyrus, and cerebellum. This finding indicates the engagement of the dorsal stream in decoding musical syntax, where auditory information is transformed to motor actions, and that this engagement is stronger among musicians than non-musicians in the presence of tonality, as discussed later.

For Western classical music perception, musicians and non-musicians achieved equally high accuracy in behavioral classification, but with a higher confidence rating in musicians, suggesting that musicians took advantage of their expertise to analyze the musical notes. Bilateral anterior superior temporal areas, bilateral left inferior frontal regions extending to bilateral precentral gyrus, insula, SMA, and cerebellum were engaged in the processing of tonal music, in line with previous studies (Koelsch et al., 2002a, 2013; Tillmann et al. 2006, Sammler et al., 2013, Farbood et al., 2015). However, whereas previous studies suggested that people perceive musical syntax implicitly, regardless of music training (Koelsch et al., 2000, Bigand et al., 2006), our results showed that music experience modulated neural activation in classical tonal music processing, though non-musicians and musicians performed equally well in behavioral classifications. Specifically, differences between musicians and non-musicians in neural activation were observed in a right-lateralized front-parieto-tempral network, covering right STG, right IFG pars triangularis and superior medial frontal gyrus, right inferior parietal gyrus, and bilateral SMA. Together with previous studies showing the role of right IFG in musical syntax processing (Cheung et al., 2018) and structural brain changes in right fronto-temporal regions linked to music training (Sato, Kirino, & Tanaka, 2015; James et al., 2014), the present findings suggest that the left fronto-temporal neural network plays an important role in musical syntactic processing in a domain-general and experience-independent way, and that the right fronto-temporal cortical areas contribute to musical syntactic processing in a musicianship-modulated way.

For impressionist music, musicians showed significantly higher accuracy in behavioral classification, as well as stronger activation in left cerebellum than non-musicians. A closer look at the neural basis among musicians and non-musicians showed that bilateral STG and bilateral IFG pars triangularis were engaged in both groups, whereas right IFG was significantly recruited only among musicians. These results suggest that the minor disruption of tonality rules in impressionist music could weaken the functions of the left IFG in resolving musical syntax. The right IFG, on the other hand, still played an important role in musical syntax processing, particularly with music training. Together with the results of classical music processing, these results indicate that music experience has an impact on the neural response to syntactic processing of tonal music – both classical tonal and impressionist (reduced tonality).

For atonal music, there were no differences between musicians and non-musicians in either neural activation or behavioral classification performance. Furthermore, atonal music could not be differentiated from random notes, either neurally or behaviorally, even among musicians. This is likely due to a lack of pitch-center relationship in atonal music, leading to an absence of structural information processing. Given that previous studies on atonal music suggested that familiarity has some effects on induced emotional responses to atonal music (Daynes, 2000), or that listeners can learn to detect or expect the avoidance of pitch repetition (Krumhansl, Sandell & Sergeant, 1987; Ockelford & Sergeant, 2012), it would be of interest for future studies to investigate whether atonal music is processed differently among musicians with more varied experiences and those with expertise in atonal music, such as the composers and conductors who have developed a positive taste for atonal music.

### 4.2 Cortical and subcortical neural networks for musical syntax processing

The IFG has been deemed to be a storage buffer required to process sequences with supra-regular structure (Fitch and Martins, 2014). Within the IFG, left pars triangularis, a part of Broca’s area, has been suggested to be involved in domain-general processing, playing a crucial role in sequence regularities, and particularly being the site of a buffer zone for syntactic computations (Sammler et al., 2011; Fitch and Martins, 2014). Previous studies have further put forward a shared resource system for domains of both language and music, seated in Broca’s area (Patel, 2003; Fedorenko et al., 2009). The role of right IFG is less clear, though some studies have suggested that the right inferior frontal area is crucial for processing specific musical syntax (Maess et al., 2001), or is sensitive to music training (Oechslin et al., 2013; Koelsch et al., 2002b). In the present study, the left pars triangularis was engaged in the syntactic processing of classical music equally for musicians and non-musicians. The right pars triangularis and pars opercularis, on the other hand, were involved to a greater extent among musicians compared to non-musicians in the syntactic processing of both classical and impressionist music. Percent signal change of different subregions of bilateral IFG further showed that right pars triangularis was sensitive to tonal differences, and that both left and right pars opercularis were sensitive to music experience differences. We therefore suggest a more precise division of labor of bilateral IFG regions in music processing: the left IFG pars triangularis carries out on-line unit relationship computations independently of music genre and music experience; the right IFG pars triangularis detects tonality and adjusts to tonal varieties, partly dependently of music experience; both left and right pars opercularis are modulated by music experience, with the right pars opercularis more dominantly so.

We also found an involvement of right anterior temporal regions, together with right frontal regions, in musical syntactic processing, especially among musicians. Furthermore, informational connectivity results revealed that higher behavioral classification accuracy among musicians was accompanied by stronger functional cooperation between right Heschl’s gyrus and right superior temporal pole. According to previous findings, temporal resolution is better in left auditory cortices, whereas spectral resolution is better in right auditory cortices (Zatorre et al., 2002). Therefore, our results suggest that right temporal regions are more engaged in musicians to achieve better performance in detecting precise changes in frequency. Together with abovementioned results on frontal regions, the present findings suggest that a right fronto-temporal network is crucial in allowing musicians to outperform non-musicians in musical syntactic processing.

The neural processing of musical syntax engages not only cortical structures but also subcortical structures, such as basal ganglia, which has been found to be activated in the processing of musical beats and music-related emotions (Frisch et al., 2003; Kung et al., 2013). In the present study, neural recruitment of pallidum and the cerebellum was found for processing tonal music in musicians. Results of the informational connectivity analysis showed that strong connectivity between the right IFG and left pallidum was positively correlated with music classification performance in musicians. Given that the sensorimotor territory of the globus pallidus internus is known to be the main output of basal ganglia, and that basal ganglia plays an important role in the storage and expression of learned sequential skills (Hikosaka et al., 2002; Doyan et al., 2009), the current finding of pallidum activation and its connection with right IFG is especially interesting. Furthermore, both globus pallidus and cerebellum are the most effective sites for deep brain simulation (DBS) in reducing motor impairments (Tewari, Fremont, & Khodakhah, 2017) and a recent study suggested that the basal ganglia and the cerebellum are interconnected at the subcortical level (Bostan & Strick, 2019). Therefore, our findings suggest that this cortico-subcortical network facilitates the perception of musical sequences, especially for the musicians, given their intensive training in music performance.

The left cerebellum was also found to be significantly more engaged in musicians compared to non-musicians in the processing of impressionist music. Among non-musicians, connectivity between the cerebellum and bilateral STG was negatively correlated with classification performance. A previous study has suggested that experience-dependent changes in cerebellum could contribute to motor sequence learning (Doyan et al., 2002), given that the motor network is important for production and perception of music (Schubotz et al. 2000), our results for the musicians suggest that the engagement of cerebellum facilitates motor sequence and musical sequence perception in turn. Further studies are needed to clarify the role of cerebellum-STG connectivity in music processing among non-musicians.

A cortico-subcortical network involving the putamen, SMA, and PMC has been proposed to be engaged in the analysis of temporal sequences and in auditory–motor interactions (Grahn & Rowe, 2009). The present study verified the engagement of these proposed regions, and in addition allowed us to have a more refined understanding of the functions of different regions. Furthermore, this cortical-subcortical connectivity is shown to be functionally correlated with behavioral performance in music genre classification and neural musical syntax processing among musicians.

### 4.3 Appreciation of tonality in music from a scientific perspective

Western classical (tonal) music has been widely appreciated due to its consonance and stability. In the present study, musicians showed stronger and more widespread neural responses to classical music compared to non-musicians. Non-musicians, though with relatively less activation than musicians, still showed stronger neural responses to classical music than to impressionist or atonal music. The higher accuracy in classifying classical musical phrases among non-musicians can be seen as evidence of implicit knowledge of musical structure even among those with minimal musical expertise. Furthermore, as described by Tonal Pitch Space (TPS) theory (Lerdahl, 1988), tension and relaxation of chords unfolding over time in classical music provide listeners with a musical context in which to generate reliable expectations.

Impressionist music, on the other hand, is well-known for feelings of ambiguity and intangibility, like impressionist paintings. This music genre places the listener in a reduced tonality context, which causes difficulty in integrating harmonics. In our study, although impressionist music and classical music both engaged similar fronto-temporal regions, they each involved specific regions as well. Furthermore, the differences between musicians and non-musicians in both behavioral and neural responses suggest that the processing of impressionist music especially involved frontal regions of the right hemisphere, and that impressionist music processing benefited from musicianship more so than classical music processing.

Lastly, the atonal genre stands opposite to tonality. Its disordered structure and unexpected musical context may well be perceived as scrambled pieces, resulting in poor performance in differentiating atonal phrases from random notes, and in a lack of significant differences in neural responses between atonal phrases and random notes, regardless of the level of music experience. There are only a few studies on tonality in neuroscience. Among them, Proverbio et al. (2015) suggested that atonal music decreased non-musicians’ heart rates and increased their blood pressure, possibly reflecting an increase in alertness and attention, and thus appeared to be perceived as being more agitating and less joyful than tonal music. The present study provides complementary results regarding the absence of “syntactic” processing in atonal music perception, and questions the “meaning” of atonal music.

Overall, by studying varying music genres and corresponding aesthetic experiences, findings in the present study allow us to gain a better understanding of neural mechanisms underlying musical syntax processing, namely how it varies across levels of tonality, and how it is modulated (or not) by music experience, and also lend strong support to music theory.

## 5 Funding

This work was supported by Art Project of National Social Science Foundation of China (No. 16BD050), National Natural Science Foundation of China (No. 31771210), National Social Science Major Project of China (No. 17ZDA323), and Science and Technology Commission of Shanghai Municipality (No. 19JC1410101).

## 6 Acknowledgments

We appreciate Shuai Wang’s suggestions and help on informational connectivity analysis.

## 7 Conflict of Interest

The authors declare that the research was conducted in the absence of any commercial or financial relationships that could be construed as a potential conflict of interest.

## 8 Data Availability Statement

The raw datasets for this study can be found in the OSF repository https://osf.io/4fejw/.

## Notes

https://osf.io/4fejw/

### Competing Interest Statement

The authors have declared no competing interest.

